# A geometric approach to human stress based on stress-related surrogate measures

**DOI:** 10.1101/688937

**Authors:** Petr Kloucek, Armin von Gunten

**Author notes:** Corresponding author: Petr Kloucek.

## Abstract

We present a predictive Geometric Stress Index (pGSI) and its relation to behavioural Entropy (*b*𝔼). *b*𝔼 is a measure of the complexity of an organism’s reactivity to stressors yielding patterns based on different behavioural and physiological variables selected as surrogate markers of stress (SMS). We present a relationship between pGSI and *b*𝔼 in terms of a power law model. This nonlinear relationship describes congruences in complexity derived from analyses of observable and measurable SMS patterns interpreted as stress. The adjective geometric refers to subdivision(s) of the domain derived from two SMS (heart rate variability and steps frequency) with respect to a positive/negative binary perceptron based on a third SMS (blood oxygenation). The presented power law allows for both quantitative and qualitative evaluations of the consequences of stress measured by pGSI. In particular, we show that elevated stress levels in terms of pGSI leads to a decrease of the *b*𝔼 of the blood oxygenation as a model of SMS.

## INTRODUCTION

This paper is an extension of our previous spectral theory of human stress, Kloucek and von Gunten (2018), providing a posteriori analysis of stress experienced by a human subject. Here, we explore the possibility to predict mental and physical stress based on a finite number of measurements using various types of artificial intelligence.

Continuous psychological stress monitoring in daily life is important. There are two conventional methods to measure psychological stress, i.e., self-report and body fluid analysis. The self-report method is hard put to monitor human stress consistently due to the lack of standards for stress status. The body fluid analysis is invasive and cannot measure stress continuously.

We intend to illustrate the potential of complexity analytics using physiological and behavioural data of a few normal subjects. More specifically, we decided to use heart frequency variability (HFV) and step frequency (SF) for the analyses as predictor variables of the oxygen saturation in the blood (SO_2_) as a binary perceptron approximated by peripheral blood oxygenation (S_p_O_2_). These variables will serve as surrogate markers of stress (SMS). We compute the heart frequency variability from HF.

The purpose of this communication is fourfold.

First, we introduce the predictive Geometric Stress Index using HF, SF and SO_2_. Complexity plays an important role in the objective indexing of SMS patterns Kloucek and von Gunten (2016), Kloucek and von Gunten (2018).

We use the adjective “geometric” to indicate that we compute separation curve(s) in the (0, 1)^2^ complexity space given the complexity projections of HFV × SF based on the respective values of the perceptron separating normoxemia domain(s) from hypooxemia domain(s).

Second, we re-define *b*𝔼 we have proposed elsewhere Kloucek, Zakharov, and von Gunten (2016). *b*𝔼 measures behavioural and/or physiological reactivity distribution of a sequence of different events represented by the complexity of a single pattern corresponding to, e.g., HFV. The concept resides with the assumption that *b*𝔼 should not be evenly, or nearly so, distributed in time. This approach is similar to the entropy concept in the physics measuring uneven distributions of energy among atoms. Increasing non-uniform energy distribution increases the entropy of “a non-organic system” while keeping its complexity high. Similar argumentation can be applied to living organisms Schrödinger (1944).

Third, we introduce the predictive Stress Resistance Index (pSRI) that is meant to quantify human resistance to various forms of stress. pSRI is based on perceptron values and their distances to the separation hyperplanes yielded by analyses of time-series of SMS.

Fourth, we propose a power law model linking pGSI with behavioral entropy applying it to time-series of SMS. We strive to predict stress in terms of pGSI in human subjects as measured through the evolution of complexity patterns, using a power law relating pGSI and *b*𝔼(.).

In short, we present a proof of concept study showing that complexity analysis of HF and SF and oxygen saturation can be used as SMS to predict human stress.

## METHODS

### Subjects

Eight subjects between thirty-five and fifty-five years, four men and four women agreed on carrying a Biovotion’s VSM (vital signes monitor) during normal work days including daily routines and sleep.

### Quantities measured

- Heart Frequency was estimated by means of a motion-compensating algorithm from pulse-induced variations of optical reflection from the skin under the sensor.
- SF Movement corresponds to the instantaneous whole-body activity of a human subject. The measurements were performed with a 3-axis accelerometer. The indicator is given by energy variations of low-passed filtered differentials of accelerometer measurements. SF was determined as the inverse of speed of movement.
- Blood Oxygenation (SO_2_) was measured using reflected red and infrared light supported by motion-compensating algorithms to estimate the ratio of hemoglobin molecules in arterial blood.

Skin perfusion and temperature were further measures, not used for the purpose of this study.

### Quantities used for the purpose of the study

We used HFV and SF for the analyses as predictor variables and SO_2_ as a binary perceptron. The choice of HFV and SF is meant to allow distinguishing increased physical activity (e.g. sport activity leading to congruent increase of HF and SF) from mental stress leading to incongruent HF increase and SF decrease. We felt the limitation to two variables was adequate mainly for two reasons. First, HF (usually its variability) is often used as an indicator of stress, and SF is a reasonable indicator for the intensity of physical activity. Second, we felt the use of only few variables was appropriate for the sake of simplicity for this proof of concept. With the purpose to introduce an element of prediction, we assume that the choice of SO_2_ as a binary perceptron is adequate. Preliminary analysis revealed that SO_2_ as measured over time showed a great variability lending itself for the purpose of complexity analyses. Furthermore, the term hypoxemia as used in this paper refers to lower levels of oxygenation relative to the mean oxygen saturation of its complexity. A further reason for the choice of the three SMS is that each of them has much bigger variance over time compared to the rest of the sensory data we had at our disposal (cf. also later).

### Analyses

We use Logical Regression (LR) Hosmer, Lemeshow, and Sturdivant (2013), Howell (2013), and Artificial Neural Networks (ANN) Kruse (2013), MacKay (2003), Ripley (1996), to obtain separation relative to normoxemia– hypoxemia boundaries in the *H*(*HFV*) × *H*(*SF*) complexity space based on SO_2_ perceptron binary values. *H*(⋅) denotes the Hurst exponent of the enclosed SMS, Mandelbrot and Van Ness (1968). The complexity product space is based on the self-similarity scaling of normally distributed SMS time-series that can be recorded over a meso-temporal time span Mörters and Peres (2010). A typical distribution is shown at Figure 1. Complexity expressed in terms of the Hurst exponent is closely related to the Hausdorff-Besicovitch dimension.

**Figure 1.**
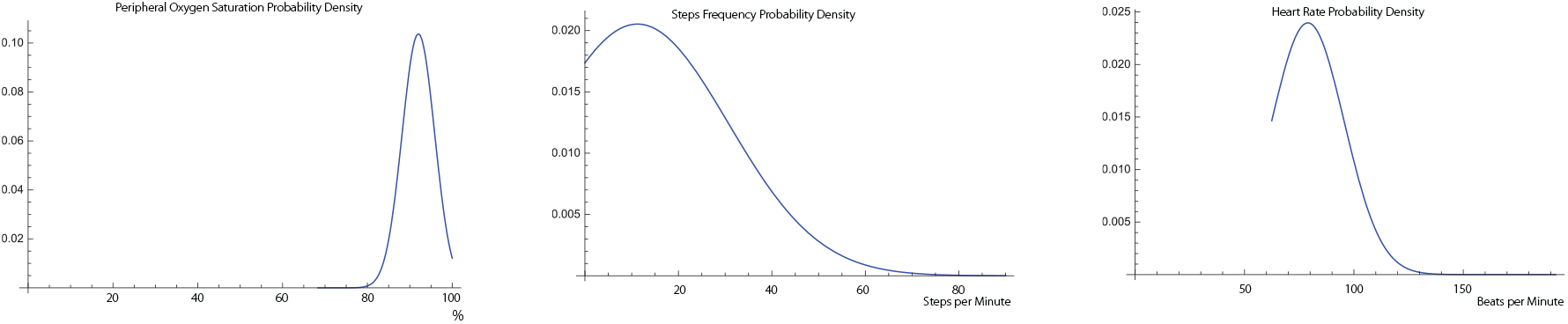
Typical probability densities of the stress indicatrix pertaining to Subject 4. The densities are computed using 1, 828 data points representing 54, 850 seconds at 30 stroboscopic resolution using histograms based on 60 bins. All three densities are very close to normal distribution and possess approximate self-similarity.

### SO_2_ as a Binary Perceptron

The crucial choice is to select which surrogate data to consider. We choose the HFV complexity and SF complexity as the two *x* and *y* axes, which express physiological (HFV) and behavioural (SF) parameters.

We consider the pGSI to depend on three surrogate time discrete processes, i.e. HFV, SF, and S_p_O_2_. The reason for the choice is that each of them have about ten to hundred times bigger variance compared to the rest of the sensory data we measured (cf. Section Variance of Some Sensory Human Data).

Furthermore, we chose SMS with high variance that also had some degree of correlation. Table 4, Table 6 and Table 7 (cf. Annex indicating correlation among HF, SF, and S_p_O_2_). The tables indicate that SO_2_ is negatively correlated with both HF and SF. The correlation tables also highlight the differences among different subjects.

Subsequently, we chose the complexity of the approximation of SO_2_ as the third surrogate data, 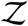. We turn this variable into a binary perceptron using formula (1). The perceptron provides a planar separation, given by smooth curve(s), of H(HFV) and H(SF). We lean on the following argument, tangentially supported by Snyder and Weathers (1977), Nikinmaa (1992), Nikinmaa and Mattsoff (1992), Nikinmaa and Jensen (1992), Chien (1970), leading to the choice of S_p_O_2_ as the binary perceptron.

A further reason for the choice of the S_p_O_2_ as binary perceptron is that stress-hormones-induced changes occur that include the *CO*_2_/*pH*-dependent decrease of the affinity of oxygen to hemoglobin due to the Bohr effect Riggs (1988), thus increasing the oxygenation potential in the tissues.

## RESULTS

### The Predictive Geometric Stress Index (pGSI)

We propose a view of some of the acquired SMS leading to the definition of pGSI.

Consider three time-discrete vectors 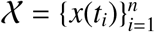, 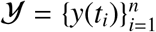, *n* ≫ 1, and 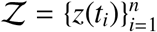 corresponding to three different sets of data representing HFV, SF and SO_2_. These quantities have different physical units and different ranges. We remove these discrepancies by projecting segmented sub-vectors on the complexity space provided by the Hurst exponent Mandelbrot and Van Ness (1968) or, equivalently, by the Hausdorff-Besicovitch dimension Peitgen, Jügen, and Saupe (1992). Using time equidistant coarse-grained segmentation 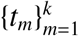, we compute

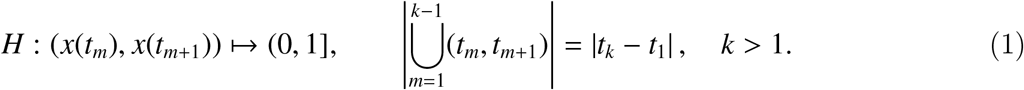

We compute such projections for all three quantities yielding coarse-grained complexity images of the three time-discrete vectors. We denote the new vectors by 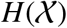, 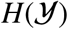 and 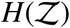, respectively. We refer to the triple 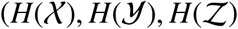 as stress indicatrix. This projection, contained in (0, 1]^3^, is not invertible for we discard micro-structural information contained in the originating time-series.

Further, we construct a binary perceptron 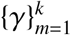 mapping 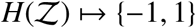 by

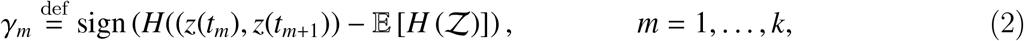

where 𝔼[.] represents the mean of the enclosed quantity.

Considering the triples 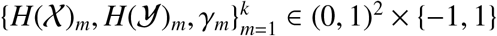 we solve an optimization problem providing “optimal”, possibly closed, curve(s) defining subdomains 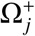 and 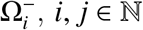, of (0, 1)^2^ such that 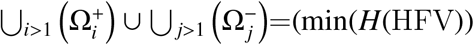, max(*H*(HFV))) × (min(*H*(SF)), max(*H*(SF))). The respective subdomains are convex hulls separating points 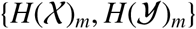 with *γ*_*m*_ = 1 from the points with *γ*_*m*_ = −1. The optimization yields the smallest number of these subdomains with the largest area at the expense of allowing a small number of opposite signs to intermix, i.e., some points with *γ*_*m*_ = −1 can appear in some 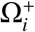. We solve this optimization problem using a combination ANN and LR. The optimization step yields a stress prediction diagram based on the complexity of the acquired SMS using geometric extrapolation that yields planar separation by a SO_2_ binary perceptron.

Finally, we define pGSI, denoted τ, by

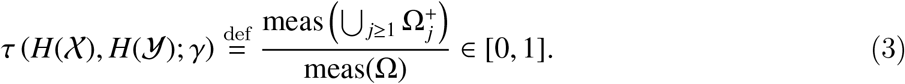

### pGSI Neutrality Baseline

We consider 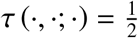 as the baseline. This approach is justified by the following observation. Let the underlining discrete time-series 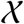, 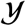 and 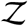 be normally distributed and self-similar. Then it is plausible to assume that

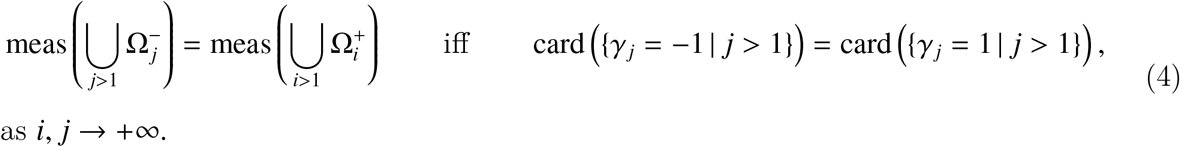

The normality assumptions are true in our computations. Normality distribution accompanied by self-similarity of the chosen surrogate markers for stress is fundamental to characterize the complexity of SMS. The examples shown in Figure 1 are computed using a histogram map with high resolution bins.

The equality represents the neutral state for it equates distribution of complexities of the surrogate data. The geometry contains more information though. The *HFV* × *SF* complexity space can be divided into four subregions reflecting complexity covariance and contra-variance with respect to higher or lower than expected individual S_p_O_2_, (c.f., Figure 2). The two covariant regions, the lower-left and upper-right quadrants, share the same short/long dynamical memories as well as negative/positive autocorrelation of either complexity of HF and SF time-series. The other two quadrants have opposite characterizations. Consider the upper-left quadrant. While the *x*−axis, representing the complexity of SF, would indicate complex SMS pattern, the HFV axis indicate a more regular pattern. These readings combined with “below-the-mean” personal SO_2_ can possibly indicate higher physical activity. Furthermore, the lower-right quadrant may indicate mental stress when the complexity of HFV and SF are reversed while the complexity of SO_2_ is still low.

**Figure 2.**
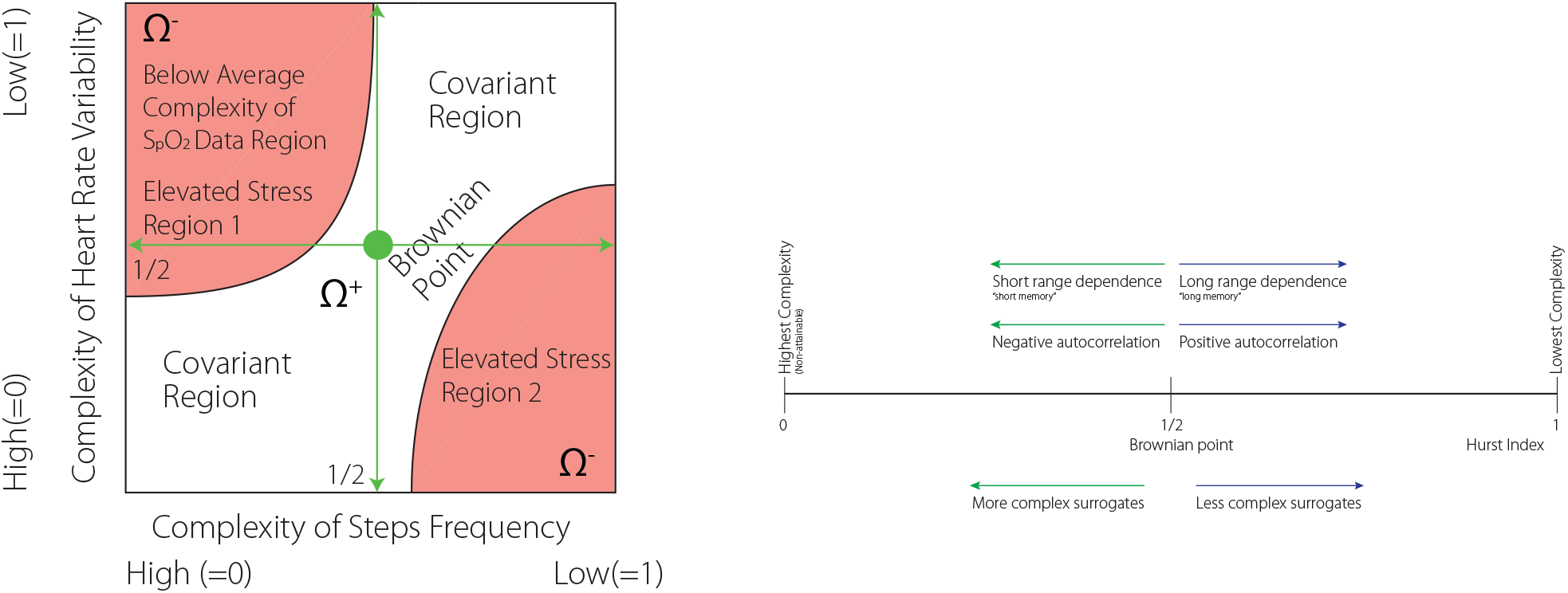
Left: Example of a segmentation of the complexity of HFV × SF space using SO_2_ as a binary perceptron. SO_2_ low complexity level, indicating possibly hypoxemia, accompanies incongruent complexity of HFV × SF combinations. Projecting the low levels of S_p_O_2_ onto the HFV × SF two-dimensional space identifies regions with undesirable HFV/SF combinations. Region 1 corresponds to physical stress (high SF): low SO_2_ complexity, high SF complexity, low HFV complexity, i.e., low S_p_O_2_ relative to individual normhypoxemia. Region 2 corresponds to mental stress: low S_p_O_2_ complexity, low SF complexity, high HFV complexity, i.e., low S_p_O_2_ relative to individual normoxemia despite low motion levels. Right: Interpretation of the complexity indices.

### Predictive Stress Resistance Index (pSRI)

The predictive Stress Resistance Index (pRSI) is a further refinement of the pGSI concept. It is based on the idea that the more data points are away from the hypoxemia – nonrmhypoxemia complexity boundary the more resistance to stress a subject will be.

The resistance index, *θ*, is defined as follows. Let

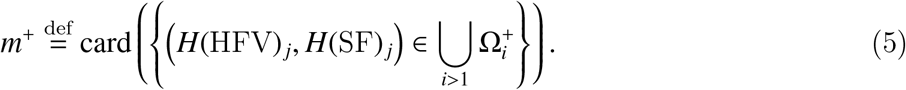

pSRI is then given by, c.f., the left drawing at Figure 5

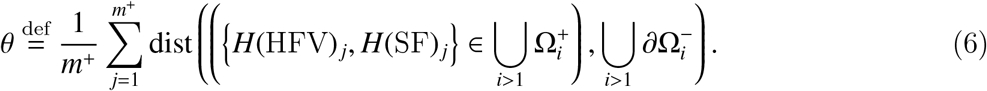

The pGSI is a global index. It can be achieved by uncountably many different configurations of the stress perceptrons. The predictive personalised stress diagram (pPSD) indicates desirable combinations of HRV/SF complexity configurations with respect to a higher level of SO_2_. The red dots correspond to *γ* = −1, the green dots correspond to *γ* = 1, i.e., to normoxemia perceptrons.

Comparing pGSI and pRSI for healthy Subject 6 (data are available upon request from authors) we conclude, as an example of the application of the pGSI/pRSI combination, that while pGSI of the Subject 6 ranks fourth, its pRSI is much lower with respect to the control group. This indicates rather medium to low ability to deal with the stress, at least if our assumptions are correct (cf. Figure 6 compared to Figure 7.

### Entropy of Behavioural Complexity

Consider a time discrete process 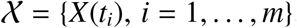, with its complexity given by the Hurst exponent, *H*(*X*(*t*_*i*_)), computed using granulation of an underlying time-series over a uniform segmentation (*t*_*i*_, *t*_*i*+1_) of (0, *T*). The Entropy of Behavioural Complexity Kloucek et al. (2016), is defined by

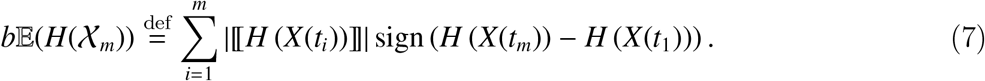

The Hurst exponent, *H*, denotes the complexity index of acquired normally distributed SMS, [[]] denotes a jump of an enclosed quantity, i.e., 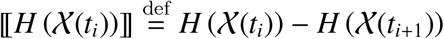, *t*_*i*_ are time equidistant points at which the function *h*: *t* ↦ *H*(*t*) has a finite jump. The signum of the difference between the complexities of previous and subsequent states indicate if the behaviours tend to a lower or higher complexity. The negative sign indicates the tendency towards higher complexity, the positive sign indicates the opposite.

We adopt the following localized time discrete notion of the entropy of behavioral complexity (7)

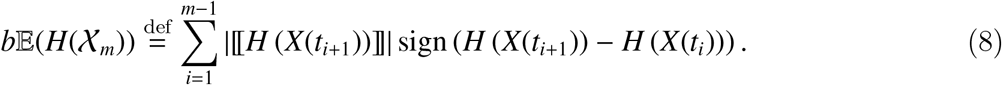

The above definition of the localized entropy is a sum of signed strengths of the complexity discontinuities.

The definition of *b*𝔼, (8), accounts also for the history of attaining certain complexity states unlike its definition (7) that accounts only for the sign of the difference between initial and terminal state.

The idea behind the (8) definition is that *b*𝔼 should be negative if the system evolves, with some probability, to a state with higher complexity and positive when the system evolves towards a lesser complexity state. Let us consider the example presented in Figure ?? using synthetic data generated by normally distributed random numbers. The red piece-wise constant function is represented by *b*𝔼 = 2.15 while the blue, decreasing function, *b*𝔼 = −2.95.

### pGSI relation to b𝔼

We propose a power law model relating pGSI to respective *b*𝔼(⋅) having the form

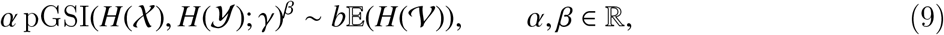

where 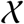, 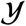 represent HFV and SF time-series. The third quantity, 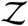, is represented by SO_2_ as the binary perceptron *γ* given by (2). The time-series 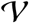 represents the remaining quantities.

We identify the power law quantities, *α* ∈ ℝ and *β* ∈ ℝ, by solving the following non-linear problem

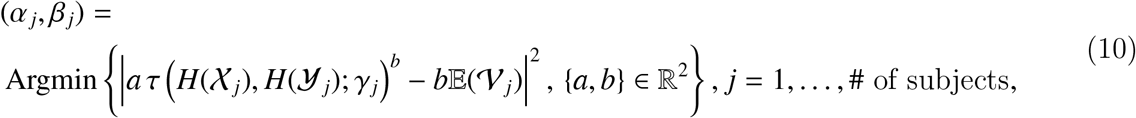

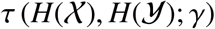 is given by (3).

The different power laws relating pGSI to *bE* of different patterns might explain the relation between stress and complexity tendencies of SMS time-series. We include the following example as an illustration. Consider 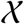 representing HF, 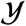 SF complexities and 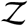 SO_2_ in the form of its binary perceptron *γ*. The computational results indicate, e.g., that

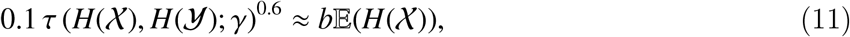

solving (10).

Scaling laws are shown in Table 2 and visualised by Figure 3.

**Figure 3.**
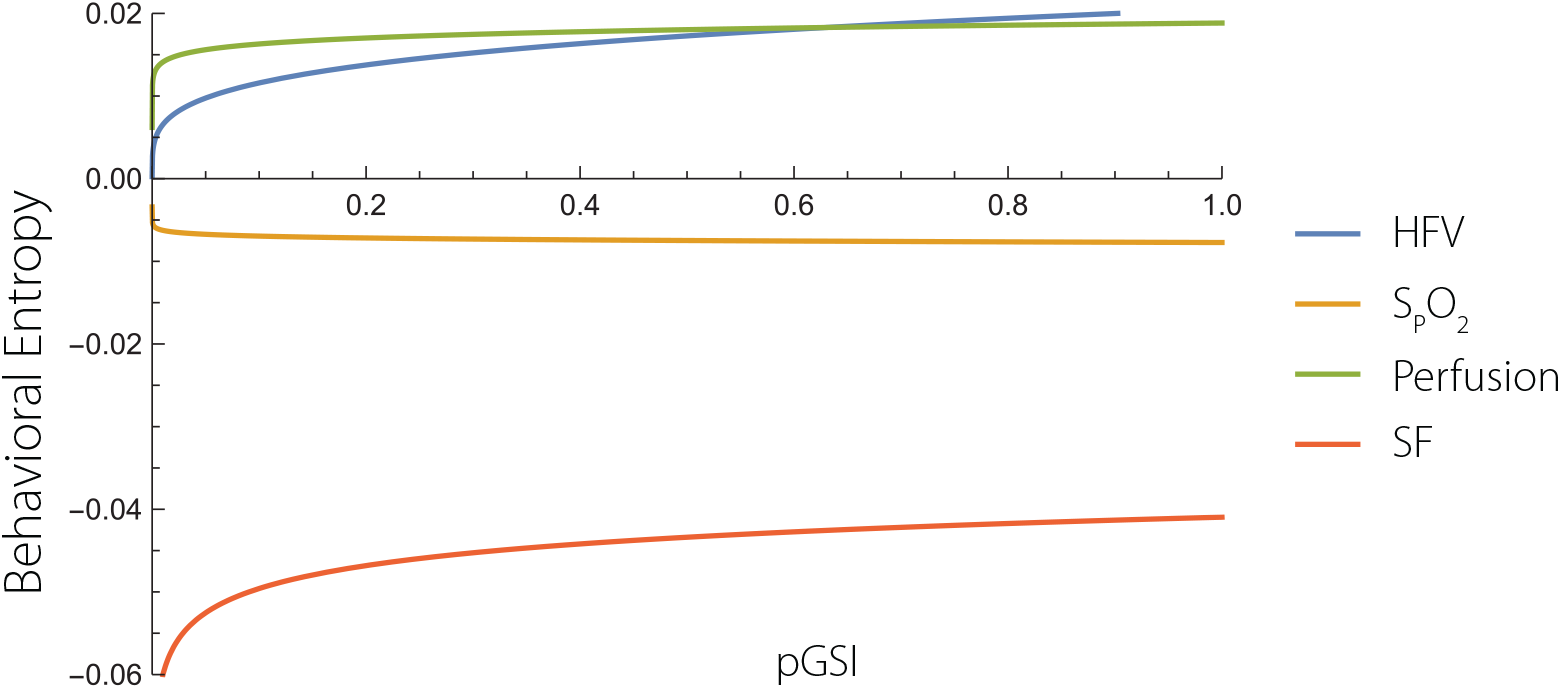
*b*𝔼 power laws. The figure is based on eight healthy subjects. The plot shows how the complexity of different SMS behave with the increasing stress index, pGSI. The curves show that complexity of all SMS increases except for the complexity of the SO_2_.

### Power Laws

Scaling laws are shown in Table 2 and visualised by Figure 3.

Figure 3 shows *b*𝔼(⋅) of HFV, SO_2_ and SF as a function of pGSI.

The curves shown at Figure 3 indicate that increased level of stress leads to increased *b*𝔼 of HFV, Perfusion and Step frequency. The only exception to this is the complexity of S_p_O_2_.

### Correlation of Heart Rate, Steps Frequency and Blood Oxygenation

We use simple Pearson product-moment correlation coefficient Pearson (1895), to estimate SMS dependency.

The data summarised by Table 4, Table 6 and Table 7 show examples of correlations among different sensory quantities of three different subjects. The first two tables correspond to healthy men and women, respectively, the third table corresponds to a female runner, during a typical working day including night/sleep readings. The first two tables show nearly equal negative correlation among HF/SO_2_, SF/SO_2_ while that third table indicates positive HF/S_p_O_2_ and nearly none SF/SO_2_ correlations.

### Mathematical Technicalities

We address in this section some subtle points related to a construction of the personal relative hypoxemia – normoxemia domain partitions in the *H*(HFV) × *H*(SF) × SO_2_ space, on which we can perform integration in order to compute domains areas to be able to compute pGSI, we can identify domains separation curves, we can compute distances to the separation boundaries, and we can decide which of the acquired SMS belong to which subdomain to be able to compute pSRI.

We use both ANN Cain (2017), Schalkoff (1997) and LR Harrell (2001), Everitt (2009), Rossi (2010), Bolstad (2010), in parallel to process the complexity indices of the acquired SMS. The reason we use two different techniques is to deal more effectively with small data sets. In the next step, we apply three different techniques to identify clusters of points forming relative hypoxemia and normoxemia subdomains, i.e., Bray-Curtis Dissimilarity measure/distance (a non-Euclidian distance) Greenacre (2017), Cutsem (1994), Krebs (1999), Chebyschev Distance Cantrell (2000), and Normalized Squared Euclidian Distance. We use Calinski-Harabasz cluster criterion, Caliński and Harabasz (1974). We select then the result with the least number of clusters. We then compute convexification of the respective clusters as a coarse-grained partitioning of the effective domain *H*(HFV) × *H*(SF) ⊂ (0, 1)^2^. We thus allow some hypoxemia points to belong to normoxemia subdomains and vice-versa. These steps fundamentally simplify subsequent construction of Delaunay triangulations Lee and Schachter (1980), Field (1988), based on the identified points in the complexity effective domain of the respective subdomains. Typically, we use 60 × 60 mesh points. Lastly, we disconnect the respective subdomains by small layers improving the quality of the triangulation by avoiding edges of the opposite classification. An example of the outcome of these procedures is shown in Figure 4.

**Figure 4.**
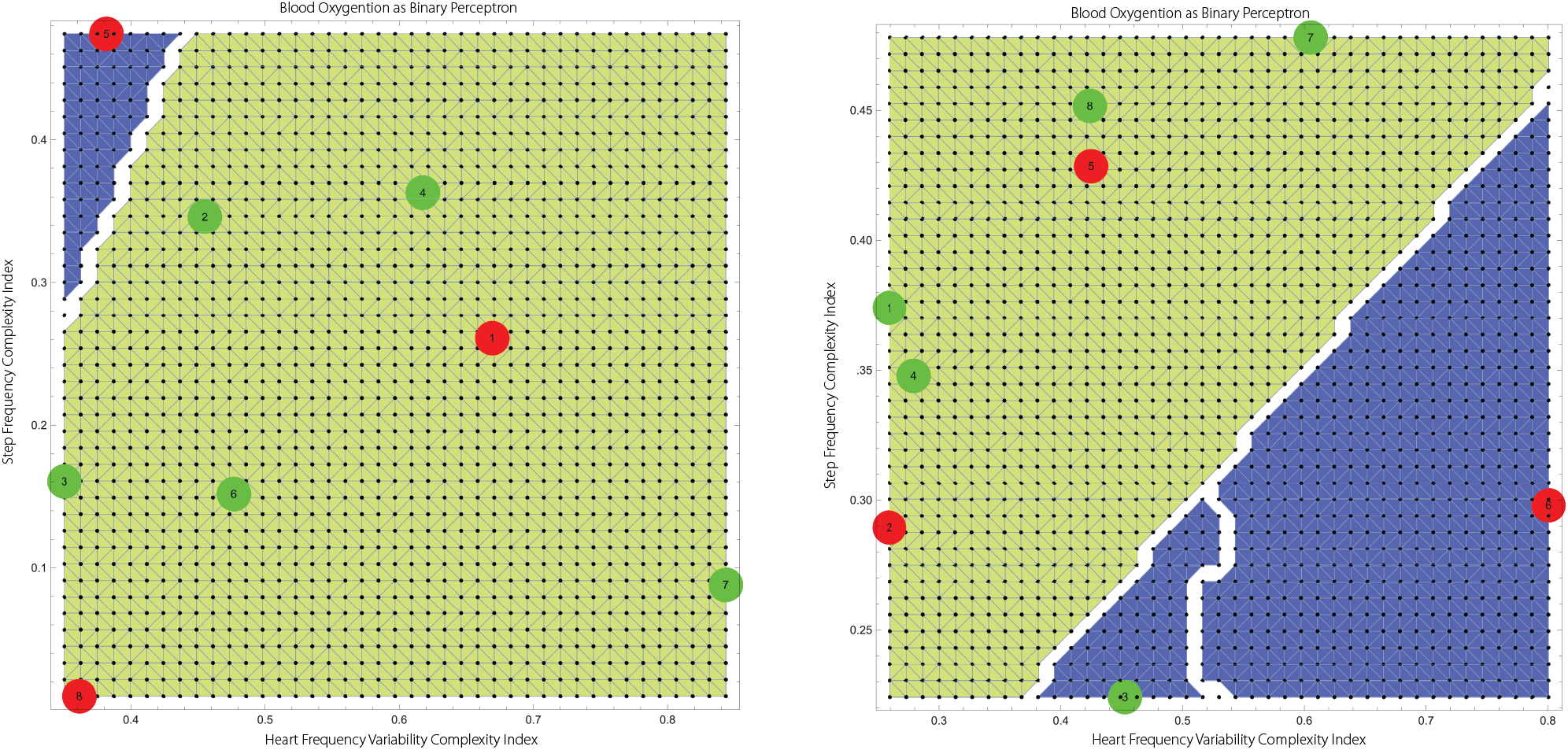
The Delaunay mesh and its separation to normoxemiaand hypoxemia subdomains of Subjects 4 and 6, respectively, used to compute both pGSI and pSRI. The predictive component of the analysis is associated with the assumption that the boundaries Γ_*i*_, *i* = 1, 2, should remain stable while the complexity data points can move around the effective complexity domain *H*(HFV) × *H*(SF).

### Results Pertaining to Human Stress and Stress Resistance

Below we report a number of findings applying our theory to real individuals. The density plot on the right at Figure 5 shows an example indicating the personal hypoxemia(blue) – normoxemia(yellow) boundary between *H*(*HFV*) and *H*(*SF*) determined by non-linear optimization using the personal SO_2_ perceptron.

**Figure 5.**
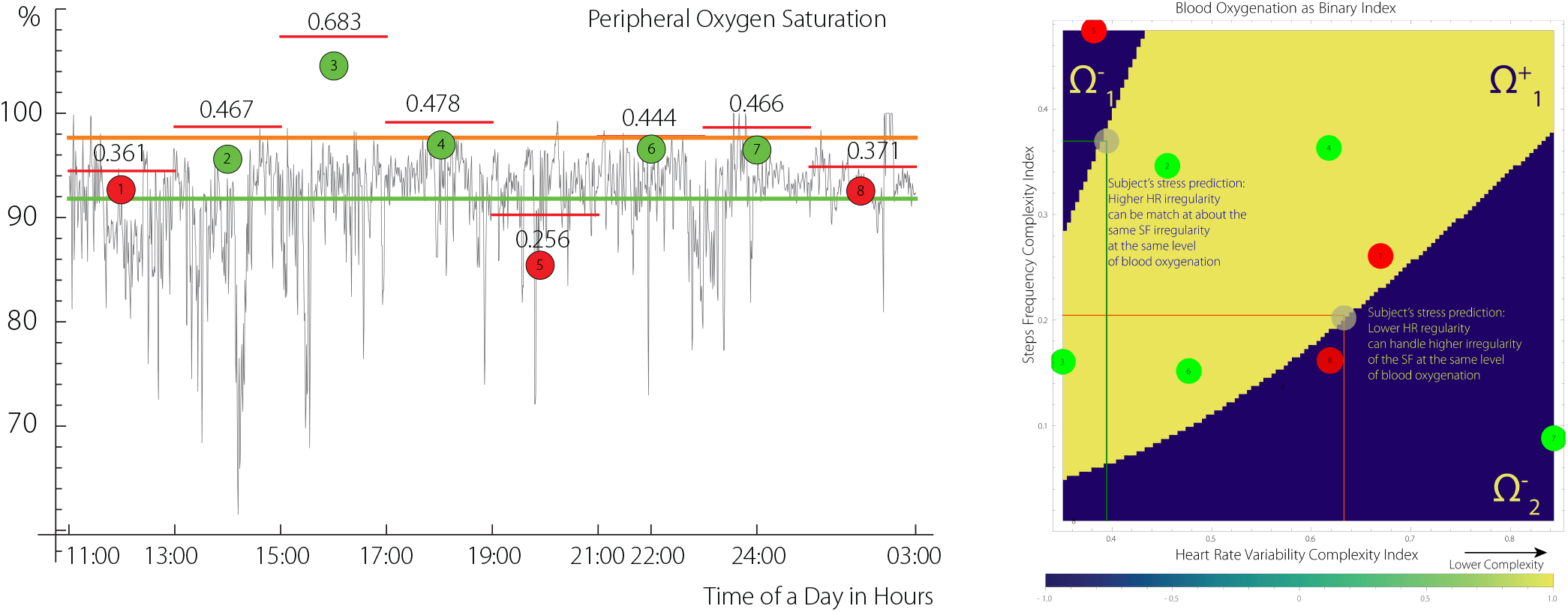
The left plot. Segmentation of SO_2_ and its projection on *H*(*HFV*) × *H*(*SF*) space in the form of SO_2_ using ANN. The step-like function indicates the value of the Hurst index for each segment. The right plot. The yellow color indicatesSO_2_-perceptron value *γ* = 1, given by (2), Ω^+^, i.e., normhypoxemia, blue colour indicates *γ* = −1, 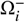, *i* = 1, 2. The grey circles are positioned at the boundary of a convex hull of certain number of points with *γ* = +1. The analyzed data correspond to a human subject encapsulating 15 hours of SMS acquisition. Each segment contains about 225 data points. The green horizontal line indicates mean of SO_2_, the orange represents the mean of the complexity segments.

To interprete the density projection shown in Figure 5, consider two different scenarios. Focusing on the lower boundary of the normoxemia – hypoxemia domain, complexity of HFV, i.e., *H*(HFV) > 1/2, exhibits lower complexity compared to SF complexity, *H*(SF) < 1/2, with a ratio of approximately 1: 3. The grey point at this boundary illustrates this scenario. The combination might represent physical activity of a trained and healthy subject.

The second scenario, represented by both *H*(HFV), *H*(SF) being below 1/2, shown by the grey dot at the upper boundary of the *γ* = 1 subdomain, indicates that HFV and SF complexities approximately match. Consequently, the first scenario might correspond to a physical stress (high activity), while the second scenario corresponds to a mental stress represented by high complexity of HFV with lower complexity of SF.

Consider segment # 8 (24:00 - 03:00) shown at Figure 5 that corresponds to a period of sleep, in which higher complexity of HFV is accompanied by a near absence of SF complexity and a lower SO_2_ complexity. Segment # 3 (15:00 - 17:00) is approximately opposite to segment # 8. The segment # 4 has all the characteristics of the first scenario, i.e. physical activity.

### pGSI and pSRI

The figure Figure 7 shows comparison between the two measures, pGSI and pSRI. Comparing pGSI and pRSI for Subject 6 we conclude, as an example of the application of pGSI/pRSI combination, that while pGSI of Subject 6 ranks fourth, its pRSI is much lower. According to our interpretation, this may indicate rather medium to low ability to deal with stress (c.f., Figure 6 compared to Figure 7).

**Figure 6.**
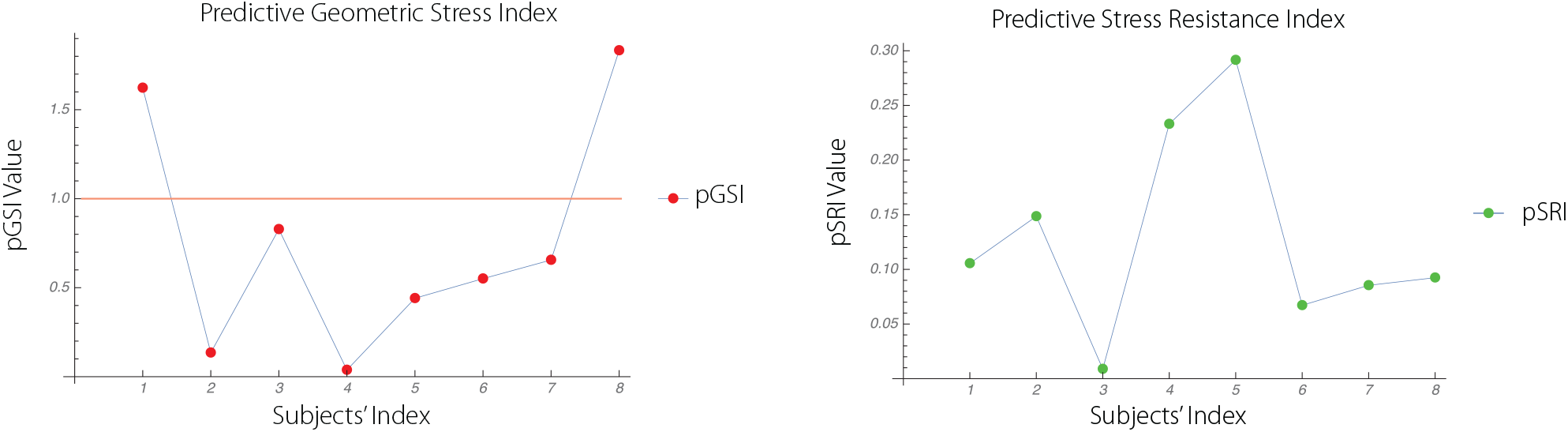
pGSIs (left) and pSRIs (right) of eight subjects. The orange horizontal separation line indicates distinction between lower and higher pGSI (c.f. Section pGSI Neutrality Baseline). Comparison of Subjects # 4 and # 5 shows that pGSI and pSRI might be also inversely related.

**Figure 7.**
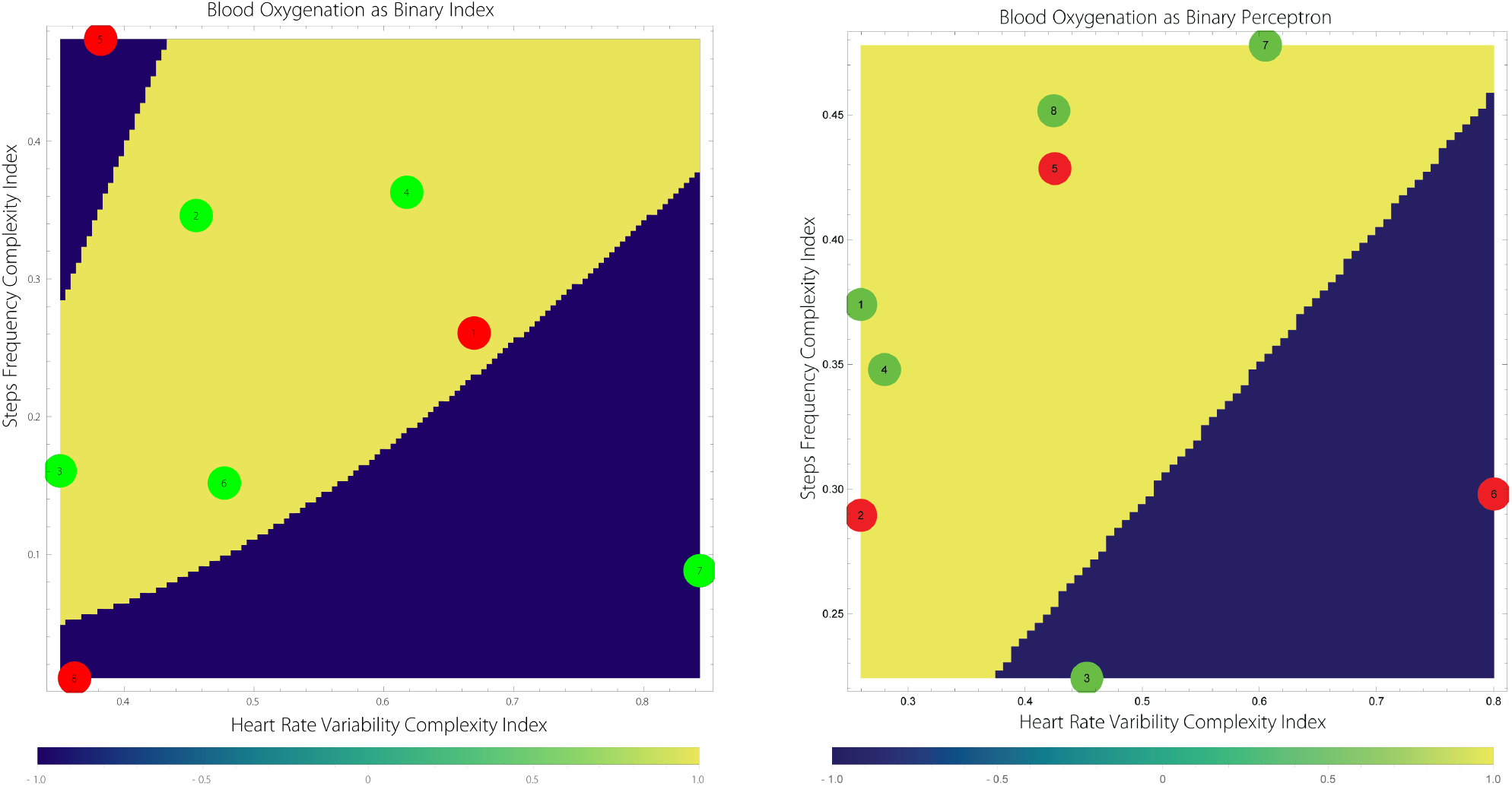
The predictive stress diagrams of Subjects 4 (left) and 6 (right) shown are generated using complexity and ANN, LR analyses.

## DISCUSSION

The main idea behind our approach is to use three dimensional phase spaces to model human stress. Our approach is based on the use of SO_2_ as the binary perceptron as well as HFV and SF as surrogate markers for SMS.

The pGSI index separates high physical activity from what we interpret as mental stress. Our analyses suggests that subjects can be distinguished regarding their overall SMS levels. Our analysis also suggests that stress is not necessarily low during sleep. Both indices, i.e., H(HFV) and H(SF) complexities correspond well with the HF and SF raw data. Low stress modes typically exhibit a positive correlation between HF and S_p_O_2_ while high stress modes have the opposite impact.

The results obtained using the geometric indices are very similar to those based on spectral theory Kloucek and von Gunten (2018). However, the spectral concept is very different from the geometric one. The combination of HF and SF complexity changes over time predicts SO_2_ complexity. Based on the variable congruency between HF and SF and the degree of SO_2_ complexity, behavioural states can be extrapolated (or predicted) as either being in the normal, high-physical activity, or mental stress realm.

For our approach using SMS to achieve clinical relevance we will have to provide evidence of correlation of the results produced by our approach with those obtained through measurements of other indicators of mental stress status. Measuring subjective stress levels or dosing stress hormones in blood or saliva such as α-amylase, cortisol or adrenalin, as well as others are necessary to prove clinical usefulness. However, none of the measures just mentioned above can be considered absolute gold standards of stress measurements. Subjective assessment of stress may be hampered in subjects with psychiatric disorders and vary widely among the normal population. Measures of hormones or neurotransmitters in blood or saliva are necessarily coarse-grained over time as they are invasive procedures and constitute no realistic approach in clinical settings. HF variability is sometimes used as another measure of stress and may be considered a gold standard for stress measures. Thus, the relationship between heart rate variability and salivary cortisol levels has been proven (Alberdi et al., 2016). However, the similarity of results of our spectral and geometric approach suggests our approach is promising.

The novelty of the proposed model of stress allows for prediction of building of a stress.

## ANNEXE

**Table 1.** *b*𝔼 of 8 subjects. *b*𝔼 is computed from equidistant time segments of acquired SMS encompassing about 15 hours of data acquisition.

| Quantities | HFV | Blood Oxygenation | Perfusion | Skin Temperature | Relative Movement | Steps Frequency |
| --- | --- | --- | --- | --- | --- | --- |
| Subject 1 | 0.000187194 | -0.0247733 | 0.0146822 | -0.087012 | -0.214842 | -0.00867333 |
| Subject 2 | 0.00470825 | 0.0169768 | 0.0169838 | -0.00218787 | -0.026098 | -0.0206288 |
| Subject 3 | -0.00816192 | -0.112449 | -0.0693975 | -0.112396 | -0.0567872 | -0.0224971 |
| Subject 4 | -0.0439746 | 0.0014295 | -0.0884581 | -0.0545995 | -0.0105887 | -0.0358108 |
| Subject 5 | 0.07338 | -0.0185053 | 0.184398 | -0.223067 | -0.0205703 | -0.2314 |
| Subject 6 | 0.0235307 | -0.00451071 | 0.0556966 | 0.0344798 | -0.0328561 | 0.0111028 |
| Subject 7 | -0.0017095 | 0.0120759 | 0.0133848 | -0.0466192 | -0.0314652 | -0.0339552 |
| Subject 8 | 0.0906186 | 0.0362499 | 0.0201563 | 0.0450308 | -0.257795 | -0.004041 |

**Table 2.** The the power law is estimated using 8 subjects. The *b*𝔼 is computed from SMS encompassing about 15 hours of data acquisition per subject.

| Behavioral Entropy/Quantities | HFV | Blood Oxygenation | Perfusion | Skin Temperature | Relative Movement | Steps Frequency |
| --- | --- | --- | --- | --- | --- | --- |
| $bE_{(HFV)}$ (Amplitude/Exponent) | 0.07/ 0.4 | | | | | |
| $bE_{(SO_2)}$ (Amplitude/Exponent) | | 0.1/0.1 | | | | |
| $bE_{(Perfusion)}$ (Amplitude/Exponent) | | | -0.0003/5. | | | |
| $bE_{(Skin Temp)}$ (Amplitude/Exponent) | | | | -0.2/-0.3 | | |
| $bE_{(Movement)}$ (Amplitude/Exponent) | | | | | -0.2/-0.3 | |
| $bE_{(SF)}$(Amplitude/Exponent) | | | | | | -0.2/0.2 |

### Correlation tables

**Table 3.** The complex stress characterization of 8 subjects. The Fatigue Recovery Index, introduced in Kloucek and von Gunten (2016), is abbreviated as FRI. *b*𝔼 is computed from equidistant two hours time segments of SMS encompassing about 15 hours of healthy human data acquisition. Comparing Subjects 4 and 6, we conclude that lower pGSI is accompanied by higher pSRI, and lower tendency complexities of HFV, SF and S_p_O_2_. The data are corroborated by fatigue recovery in the *Var*(HF) × *H*(SF) space with the ratio of about 1: 30 in favor of the highly trained Subject 4. Also the posteriori indices, CSSIs, are consistent with the two other indicators. Consider the subject #4: the lowest predictive exposer to stress, second highest resistance to stress, second lowest posteriori stress level during a working day cycle including sleep, second lowest FRI, all the three Behavioral Entropies indicating higher reactivitivness with respect to HFV, SF and S_p_O_2_ Kloucek and von Gunten (2018).

| Quantities | pGSI | pSRI | CSSI | $bE(HFV)$ | $bE(SF)$ | $bE(S_pO_2)$ | FRI |
| --- | --- | --- | --- | --- | --- | --- | --- |
| Subject 1 | 1.44898 | 0.118618 | 5.08567 | -0.094372 | -0.214842 | -0.463229 | 5.08567 |
| Subject 2 | 0.0936795 | 0.156836 | 3.65345 | -0.0669521 | -0.203418 | 0.278933 | 13.3114 |
| Subject 3 | 3.90593 | 0.0201643 | 3.83594 | -0.207423 | -0.0567872 | -0.0693975 | 4.49745 |
| Subject 4 | 0.0356675 | 0.224131 | 3.40091 | -0.100361 | -0.596833 | -0.131314 | 4.30004 |
| Subject 5 | 0.485007 | 0.254013 | 4.6757 | -0.308734 | -0.135028 | 0.184398 | 3.46533 |
| Subject 6 | 0.5625 | 0.14543 | 3.95665 | 0.183051 | -0.447905 | 0.113521 | 136.667 |
| Subject 7 | 1.23153 | 0.075056 | 3.1995 | -0.0466192 | -0.0813598 | 0.153274 | 8.24744 |
| Subject 8 | 1.6087 | 0.11549 | 7.32154 | 0.0450308 | -0.36093 | 0.0201563 | 10.5815 |

### Two variance tables

**Table 4.**
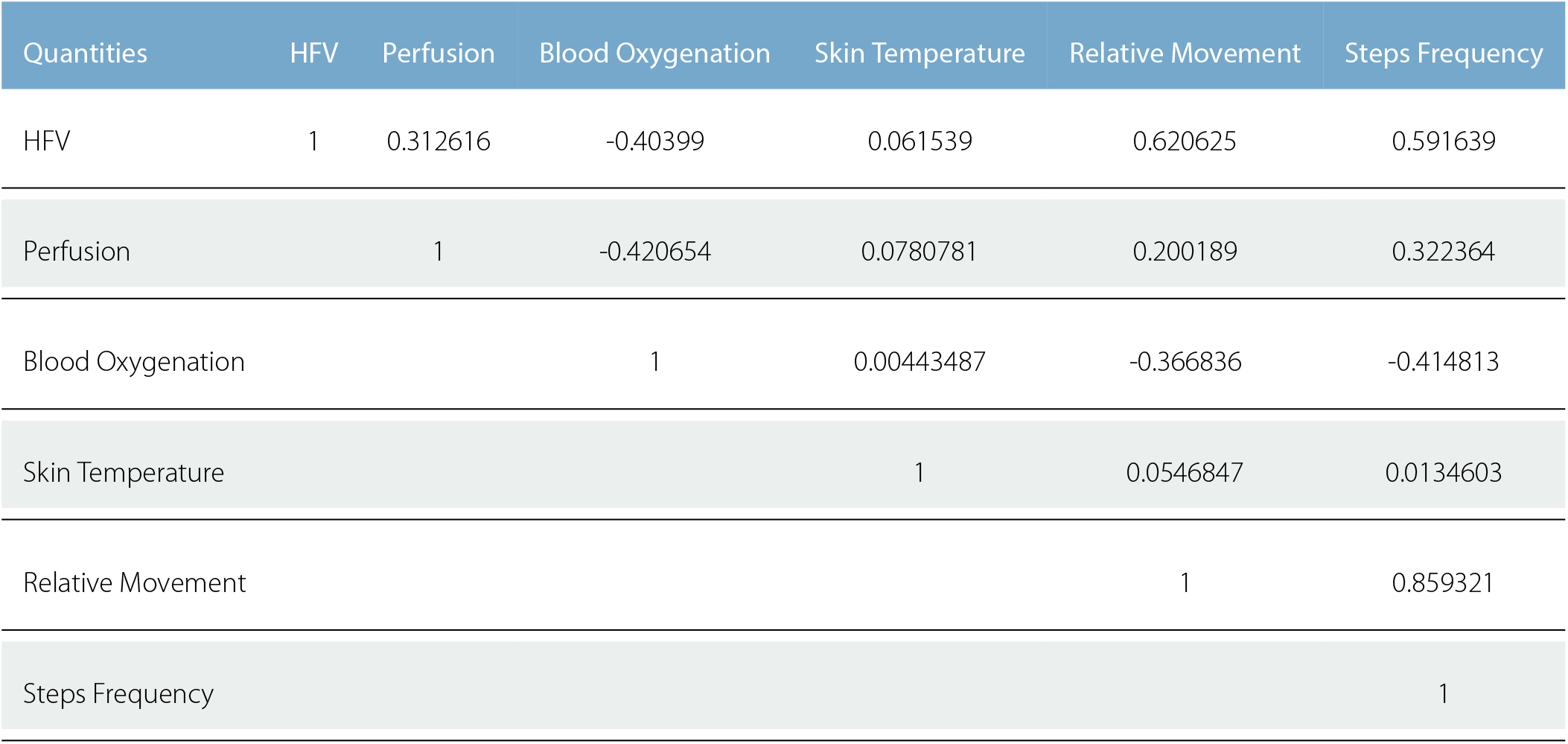
The correlation of the SMS for Subject 1. The correlation was obtained from 894 equidistant time segments of acquired data.

| Quantities | HFV | Perfusion | Blood Oxygenation | Skin Temperature | Relative Movement | Steps Frequency |
| --- | --- | --- | --- | --- | --- | --- |
| HFV | 1 | 0.312616 | -0.40399 | 0.061539 | 0.620625 | 0.591639 |
| Perfusion |  | 1 | -0.420654 | 0.0780781 | 0.200189 | 0.322364 |
| Blood Oxygenation |  |  | 1 | 0.00443487 | -0.366836 | -0.414813 |
| Skin Temperature |  |  |  | 1 | 0.0546847 | 0.0134603 |
| Relative Movement |  |  |  |  | 1 | 0.859321 |
| Steps Frequency |  |  |  |  |  | 1 |

**Table 5.** The correlation of the SMS for subject 2. The correlation was obtained from 431 equidistant time segments of acquired data.

| Quantities | HFV | Perfusion | Blood Oxygenation | Skin Temperature | Relative Movement | Steps Frequency |
| --- | --- | --- | --- | --- | --- | --- |
| HFV | 1 | -0.0144155 | -0.281479 | -0.178694 | 0.445759 | 0.366347 |
| Perfusion |  | 1 | -0.0231772 | -0.090395 | 0.103687 | 0.0946516 |
| Blood Oxygenation |  |  | 1 | 0.121536 | -0.275235 | -0.268467 |
| Skin Temperature |  |  |  | 1 | -0.337818 | -0.254233 |
| Relative Movement |  |  |  |  | 1 | 0.797544 |
| Steps Frequency |  |  |  |  |  | 1 |

**Table 6.**
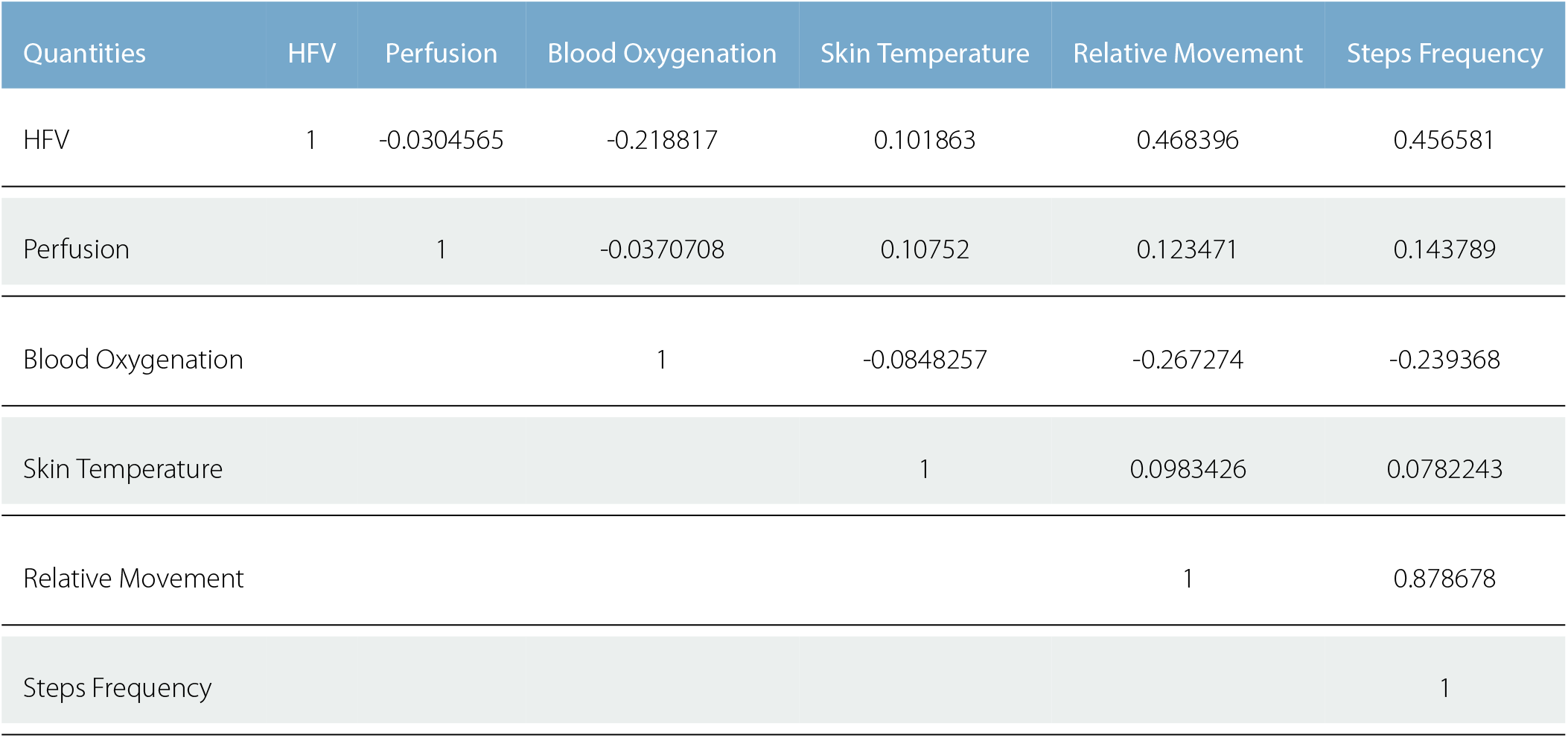
The correlation of the SMS for subject 3. The correlation was obtained from 1160 equidistant time segments.

**Table 7.** The correlation of the SMS for Subject 4. The correlation was obtained from 374 equidistant time segments.

| Quantities | HFV | Perfusion | Blood Oxygenation | Skin Temperature | Relative Movement | Steps Frequency |
| --- | --- | --- | --- | --- | --- | --- |
| HFV | 1 | -0.17864 | 0.251414 | 0.0637194 | 0.625233 | 0.676598 |
| Perfusion |  | 1 | -0.413867 | 0.0450275 | 0.124743 | 0.0341549 |
| Blood Oxygenation |  |  | 1 | -0.0803077 | -0.107687 | -0.0287573 |
| Skin Temperature |  |  |  | 1 | -0.0155391 | 0.0199515 |
| Relative Movement |  |  |  |  | 1 | 0.863616 |
| Steps Frequency |  |  |  |  |  | 1 |

**Table 8.** The correlation of the SMS for subject 5. The correlation was obtained from 1826 equidistant time segments.

| Quantities | HFV | Perfusion | Blood Oxygenation | Skin Temperature | Relative Movement | Steps Frequency |
| --- | --- | --- | --- | --- | --- | --- |
| HFV | 1 | -0.25438 | -0.286885 | -0.303626 | 0.699491 | 0.580695 |
| Perfusion |  | 1 | 0.141447 | 0.422901 | -0.044777 | -0.014162 |
| Blood Oxygenation |  |  | 1 | 0.102708 | -0.148153 | -0.041951 |
| Skin Temperature |  |  |  | 1 | -0.055175 | 0.0165696 |
| Relative Movement |  |  |  |  | 1 | 0.905904 |
| Steps Frequency |  |  |  |  |  | 1 |

**Table 9.** The correlation of the SMS for subject 6. The correlation was obtained from 642 equidistant time segments.

| Quantities | HFV | Perfusion | Blood Oxygenation | Skin Temperature | Relative Movement | Steps Frequency |
| --- | --- | --- | --- | --- | --- | --- |
| HFV | 1 | 0.10924 | -0.129858 | 0.0153075 | 0.350668 | 0.267524 |
| Perfusion |  | 1 | -0.146234 | 0.112445 | 0.229024 | 0.206663 |
| Blood Oxygenation |  |  | 1 | -0.187969 | -0.311809 | -0.207137 |
| Skin Temperature |  |  |  | 1 | -0.0353194 | -0.0188643 |
| Relative Movement |  |  |  |  | 1 | 0.778953 |
| Steps Frequency |  |  |  |  |  | 1 |

**Table 10.** The correlation of the SMS for subject 7. The correlation was obtained from 1658 equidistant time segments.

| Quantities | HFV | Perfusion | Blood Oxygenation | Skin Temperature | Relative Movement | Steps Frequency |
| --- | --- | --- | --- | --- | --- | --- |
| HFV | 1 | 0.167969 | -0.36686 | -0.0750689 | 0.630799 | 0.608615 |
| Perfusion |  | 1 | -0.253866 | 0.200398 | 0.284085 | 0.311865 |
| Blood Oxygenation |  |  | 1 | 0.217739 | -0.437872 | -0.389442 |
| Skin Temperature |  |  |  | 1 | -0.0674336 | -0.00604729 |
| Relative Movement |  |  |  |  | 1 | 0.93125 |
| Steps Frequency |  |  |  |  |  | 1 |

**Table 11.** The correlation of the SMS for subject 8. The correlation was obtained from 452 equidistant time segments.

| Quantities | HFV | Perfusion | Blood Oxygenation | Skin Temperature | Relative Movement | Steps Frequency |
| --- | --- | --- | --- | --- | --- | --- |
| HFV | 1 | -0.0450726 | -0.112492 | -0.107827 | 0.483597 | 0.479533 |
| Perfusion |  | 1 | -0.107982 | -0.0231544 | -0.0138105 | -0.0816 |
| Blood Oxygenation |  |  | 1 | 0.130734 | -0.236126 | -0.170944 |
| Skin Temperature |  |  |  | 1 | -0.0825037 | -0.032942 |
| Relative Movement |  |  |  |  | 1 | 0.901963 |
| Steps Frequency |  |  |  |  |  | 1 |

**Table 12.** The variance of the SMS for Subject 4. The variance was obtained from 374 equidistant time segments of acquired data.

| Quantities | Segment 1 | Segment 2 | Segment 3 | Segment 4 | Segment 5 | Segment 6 | Segment 7 | Segment 8 |
| --- | --- | --- | --- | --- | --- | --- | --- | --- |
| HFV | 22.3652 | 37.8688 | 43.3020 | 70.4190 | 66.9771 | 20.2466 | 27.4566 | 27.9333 |
| Perfusion | 0.00210023 | 0.00303487 | 0.0142073 | 0.00308452 | 0.00448286 | 0.00580235 | 0.0437197 | 0.00561481 |
| Blood Oxygenation | 16.4855 | 44.9826 | 34.3267 | 12.5918 | 12.6589 | 19.9398 | 7.82296 | 4.68279 |
| Skin Temperature | 0.915423 | 0.0705794 | 0.018323 | 0.0235593 | 0.0229517 | 0.0999276 | 0.377792 | 0.27331 |
| Relative Movement | 0.522469 | 0.51333 | 1.06422 | 3.57759 | 1.56432 | 0.498561 | 0.156739 | 0.0188911 |
| Steps Frequency | 54.9058 | 30.7965 | 102.836 | 613.618 | 333.916 | 22.4165 | 3.5368 | 0.00266509 |

**Table 13.** The variance of the SMS for Subject 6. The variance was obtained from 642 equidistant time segments of acquired data encompassing about 15 hours of data acquisition.

| Quantities | Segment 1 | Segment 2 | Segment 3 | Segment 4 | Segment 5 | Segment 6 | Segment 7 | Segment 8 |
| --- | --- | --- | --- | --- | --- | --- | --- | --- |
| HFV | 339.983 | 101.478 | 118.456 | 146.107 | 522.311 | 34.4726 | 19.3977 | 13.9784 |
| Perfusion | 0.0543121 | 0.0315185 | 0.00918733 | 0.00217061 | 0.0114509 | 0.00109113 | 0.00178169 | 0.0134619 |
| Blood Oxygenation | 66.8304 | 68.556 | 16.1578 | 13.3593 | 12.07 | 2.83156 | 3.41918 | 8.13148 |
| Skin Temperature | 0.425757 | 0.44536 | 0.488427 | 0.197185 | 2.53583 | 0.2535 | 0.350932 | 0.0978087 |
| Relative Movement | 4.34773 | 0.263877 | 3.22692 | 2.39485 | 5.16624 | 0.251206 | 0.31262 | 0.712321 |
| Steps Frequency | 447.909 | 20.6665 | 356.858 | 216.062 | 772.406 | 4.34818 | 10.0686 | 22.1422 |

## DECLARATIONS

### Authors’ Contributions

Both authors contributed equally to the presented research.

### Ethics

The investigation was carried under ethics application “Indexation mathématique de mesures physiologiques multiples non-invasives en milieu réel chez des sujets sains”, CHUV, Lausanne Switzerland.

In addition each subject signed “Informed Consent” prior to measurements of the data.

The collection and handling of data has been carried out in accordance to EU current regulations, GDPR.

